# Direct analytical estimation of thermodynamic parameters of thermo-TRP channels

**DOI:** 10.64898/2026.05.21.726968

**Authors:** Mira Stoll, Michael Mazar, Inti Zumeta-Dubé, Karel Talavera

## Abstract

A subset of Transient Receptor Potential (TRP) channels display very steep temperature dependences and play key roles in thermosensation. To characterize the properties of these ‘thermoTRP’ channels, two-state close-open gating models were developed for TRPM8, TRPV1, TRPM4, TRPM5, TRPA1 and TRPM3. In this study, we met the recurrent challenge of finding an initial set of model parameters enabling effective convergence during data fitting procedures. We performed algebraic calculations to derive equations for all gating model parameters as functions of key features of thermoTRP channel data obtained from patch-clamp experiments. We used a minimal set of experimental data: the steady-state open probability and time constant of current relaxation as functions of the membrane potential determined at two temperatures. Specifically, we could express the electric distance of the gating charge and the enthalpy and entropy changes associated with the gating transitions, as functions of the voltages for half-maximal activation, the voltages for maximal time constant of current relaxation and the maximal time constant. Our results provide a method to analytically estimate an initial set thermoTRP thermodynamic parameters enabling robust subsequent nonlinear global data fitting. This approach facilitates quantitative analysis of channel thermodynamics, and has potential applications to more complex gating models, and to the study of permeation, block and other ion channel gating mechanisms.

## Introduction

The Transient Receptor Potential (TRP) proteins constitute a large and diverse group of cation-permeable channels that are activated by numerous physical and chemical cues [16]. Several of them, known as thermoTRPs, exhibit gating properties with exceptional sensitivity to thermal stimuli, and play key roles in thermosensation [17,26,15,23,19]. The quantitative characterization of thermoTRP channel behavior has been central to both the dissection of their physiological implications in thermo-sensory processes and to the elucidation of the molecular basis of their pronounced temperature sensitivity. Indeed, this has served to discern how much a given thermoTRP channel mediates the sensing of a specific temperature range at cellular and behavioral levels [17,26,15], and allows for comparisons between wild type and mutant channels in structure-function studies aimed at identifying protein motifs implicated in the responses to thermal stimuli (e.g., [25,3,8,18,7,6,22,2,9,10,28,5,29].

The thermal responses of these thermoTRPs were initially characterized by two simple parameters: the thermal threshold of activation (*T*_*thr*_) and the *Q*_*10*_ [23]. *T*_*thr*_ refers to the temperature at which TRP channel currents showed an apparent increase above background currents, while *Q*_*10*_ indicates the fold-increase in current amplitude for a 10 °C temperature change. However, as thoroughly discussed before both concepts have serious limitations [20,23]. For instance, in its original definition *T*_*thr*_ is biophysically meaningless, since thermoTRP channels do not activate at a specific temperature, but undergo smooth changes in open probability all along the physiological temperature range [1,14,21,24,27,23]. Moreover, the method used to determine *T*_*thr*_ is not precise, as it depends on the size of the background and leak currents present in each recorded cell. The *Q*_*10*_ can be misleading as well, as it depends on voltage, temperature and the size of background currents, and does not have a clear physical interpretation [20,23].

A more comprehensive way to characterize thermoTRP channel properties has been sought through the formulation of gating models based on kinetic theory [23], following the principles of ion channel biophysics research started by Hodgkin and Huxley [12] and further developed by many others [11]. Several biophysical models have been proposed to account for thermoTRP properties: i) a two-state (closed-open) model in which the rate constants of channel opening and closing depend on temperature and voltage [24,21,14,27], ii) an allosteric multi-state model considering separate temperature- and voltage-dependent transitions [1], iii) models considering that channel transitions are associated with changes in heat capacity [4,30], and iv) combinations thereof [13].

However, although clearly superior to the analysis of *T*_*thr*_ and *Q*_*10*_, the use of gating models comes with the difficulty of a more intensive data processing. Indeed, obtaining the thermodynamic model of a given channel entails determining all parameters through a fitting procedure that is rather cumbersome. The most salient hurdle is finding an initial set of model parameters that ensures proper convergence during the iterative fitting procedure. Convinced about the heuristic value of gating models, in this study we addressed this issue by deriving equations allowing to estimate all gating model parameters as functions of salient features of thermoTRP channel data obtained in patch-clamp experiments.

As a study case, we used the two-state close-open model initially developed for TRPM8 and TRPV1 using an Arrhenius rate formulation [24], and soon after for TRPM4, TRPM5, TRPM8 and TRPV1 using the more appropriate Eyring theory of absolute reaction rates [21]. We chose this model for several reasons: 1) it is the simplest one able to capture the most salient features of the thermal responses of thermoTRP channels and therefore the one for which the parameter interpretation is most straightforward, 2) it is the only gating model available accounting for channel kinetics (time dependence of the open probability), and 3) unlike any other model, it has been successfully applied to multiple thermoTRP channels, including TRPV1, TRPM8, TRPM4, TRPM5, TRPA1 and TRPM3 [24,21,14,27].

In the Eyring’s reaction rate formalism the voltage- and temperature-dependent opening and closing rate constants *α* and *β* are expressed as:

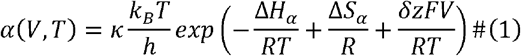

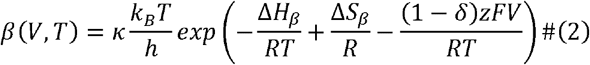

where κ is a transmission coefficient that is taken as 1, *k*_*B*_ is Boltzmann’s constant, *T* is the absolute temperature, *h* is Planck’s constant, *R* is the gas constant, Δ*H*_*α*_, *&S*_*α*_, Δ*H*_*β*_, and Δ*S*_*β*_ are the molar enthalpy and entropy changes associated with the opening and closing transitions, respectively, *F* is the Faraday constant, *V* is the membrane potential, *z* is the valence of the apparent gating charge, and *S* is the electrical coupling between the gating charge and the membrane potential. In practice, the magnitudes that can be determined experimentally are the steady-state open probability, *P*_*o*_, and the time constant of current relaxation, *τ*, as functions of temperature and voltage [24,21,14,27], These are given by:

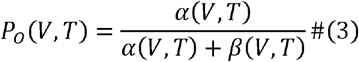

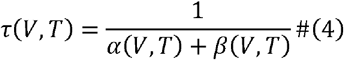

or

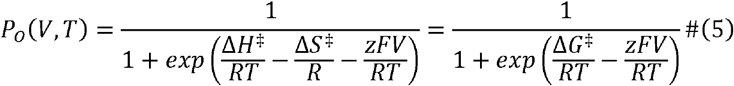

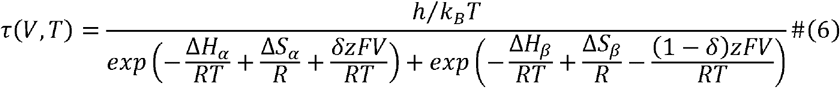

where Δ*H*^‡^ = Δ*H*_*α*_ *—* Δ*H*_*β*_, Δ.*S*^‡^ *=* Δ.*S*_*α*_ *—* Δ*S*_*β*_ and Δ*G*^‡^ *=* Δ*H*^‡^ *— T*Δ.*S*^‡^ are the differences of molar enthalpy, entropy and free Gibbs energy between the open and the closed states, respectively. In the simplest form of the model all parameters (*z, δ*, Δ*H*_*α*_, Δ*S*_*α*_, Δ*H*_*β*_, and Δ.*S*_*β*_) are temperature independent, an assumption that seems appropriate according to previous studies [24,21,14,27].

In principle, all model parameters can be determined by fitting the equation for *τ (V, T)* to the experimental voltage dependences of *τ* obtained at 2 different temperatures (i.e., *τ (V, T*_*1*_*) and τ (V, T*_*2*_*)’)*. However, fitting a 6-parameter function to a typically limited experimental data set can result in very weak constraining and high interdependence of the parameters. Note that typically there is a lack of experimental data for *τ (V,T)* due to the small current amplitudes around the reversal potential (≈ 0 mV). This limitation can be circumvented by using also the *P*_*0*_*(V,T)* set of data obtained from tail currents during the fitting procedure [24,21,14,27].

## Results

As indicated above, a successful fitting procedure requires having an initial set of model parameters that ensures convergence. In the following, we derive equations for all gating model parameters (*z, δ*, Δ*H*^‡^, Δ*S*^‡^, Δ*H*_*α*_, Δ*S*_*α*_, Δ*H*_*β*_ and Δ*S*_*β*_) as functions of salient features of thermoTRP channel data obtained in patch-clamp experiments.

### Derivation of *z*

At a given temperature, the maximal sensitivity of the open probability to voltage is obtained from the condition of maximal *∂P*_*0*_*/∂V*, or equivalently from the zero of the second derivative *∂*^2^ *P*_*o*_*/∂V*^2^.

The calculation of the first derivative yields:

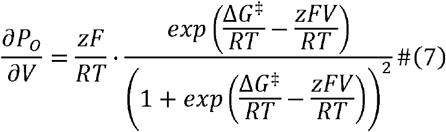

or:

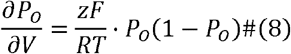

The second derivative is given by:

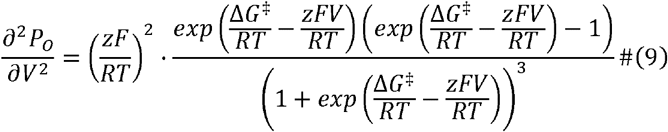

or in a more compact manner:

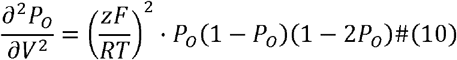

The meaningful zero of the right-hand side is obtained for *P*_*o*_ = 1/2.

The *P*_*o*_ = 1/2 condition occurs at the so-called voltage for half-maximal activation, *V*_1/2_(*T*), which is a crucial reference parameter that can be read directly from the experimental *P*_*0*_*(V,T)* curves.

Evaluating Eq. 8 in this condition yields:

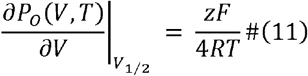

from which:

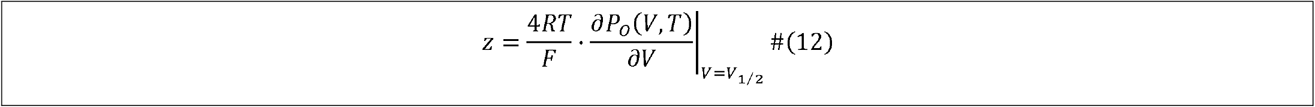

Thus, the apparent gating valence can be determined from the slope of the activation curve at the voltage for half-maximal activation. Geometrically speaking *z* can be determined from the maximal slope of the activation curve.

*V*_1/2_ (*T*) can be derived from the condition *P*_*0*_*(V,T)* = 1/2, which implies:

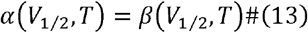

which yields that:

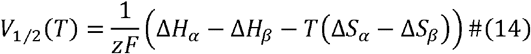

or

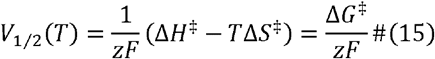

One can determine *z* also by fitting the activation curve *P*_*o*_ *(V, T’)*, with the more traditionally used Boltzmann type of equation that is written in terms of *V*_1/2_ (*T*) and the slope factor *s (T):*

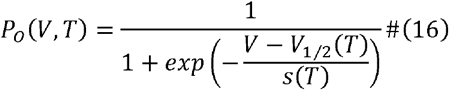

By considering that:

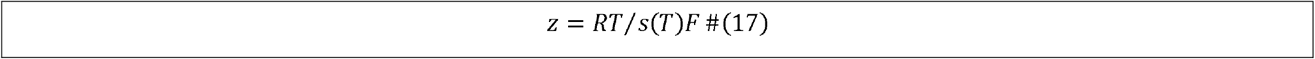

Finally, note that the slope factor is given by:

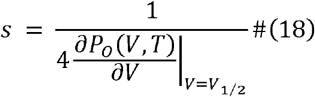

### Derivation of *δ*

The voltage dependence of the relaxation time constant shows an absolute maximum, and the voltage at which this occurs, *V*_*τmax*_, can be determined from the condition:

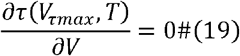

Differentiating the equation for *τ* (Eq. 4) yields:

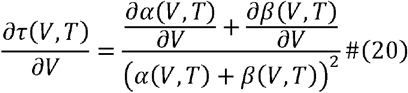

Hence, the condition for a null derivative becomes:

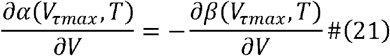

Considering Eqs. 1 and 2, these voltage derivatives are given by:

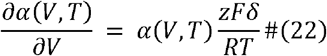

and

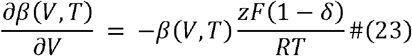

This implies

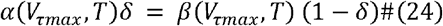

Solving for *δ*yields:

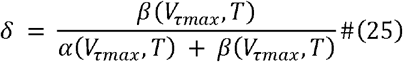

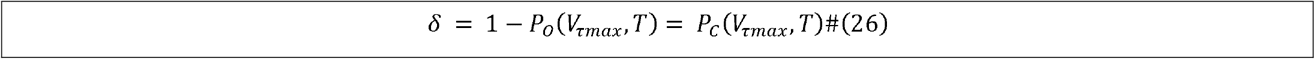

Then, using the equation for *P*_*c*_ to evaluate it in *V = V*_*τmax*_, one obtains:

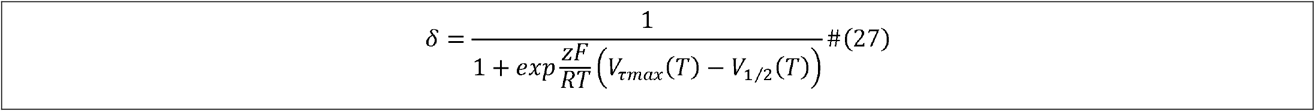

Thus, the parameter *δ*can be determined operationally from the patch-clamp data in two ways. One, using Eq. 26 and determining the closed probability at the voltage for maximal relaxation time, *V*_*τmax*_, and two, using Eq. 27, from the difference between *V*_*τmax*_ and *V*_l/2_, at any given temperature.

### Derivation of Δ*S*^‡^ and Δ*H*^‡^

In principle, Δ*S*^‡^ and Δ*H*^‡^ can be derived from the slope and the intercept of the linear fitting of Eq. 15 to the experimental dependency of *V*_1/2_ vs. *T*, respectively ([21]). However, using as a first approach one could use the experimental *V*_1/2_ values measured at just two temperatures, *T*_1_ and *T*_2_. In this case:

At temperature *T*_1_ :

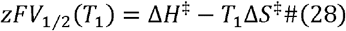

For *T*_*2*_:

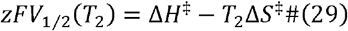

Subtracting the first from the second equation:

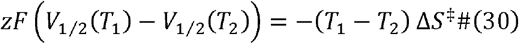

Solving for the entropy gives:

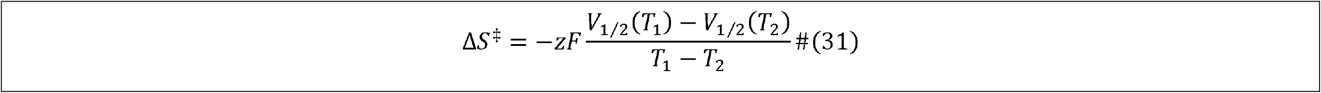

To determine the enthalpy, we multiply Eqs. 28 and 29 by *T*_2_ and *T*_1_, respectively:

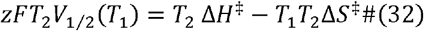

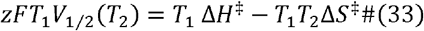

Subtracting Eq, 33 from Eq. 32 eliminates *T*_1_*T*_2_ Δ*S*^‡^:

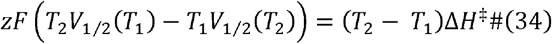

Solving for the entropy gives:

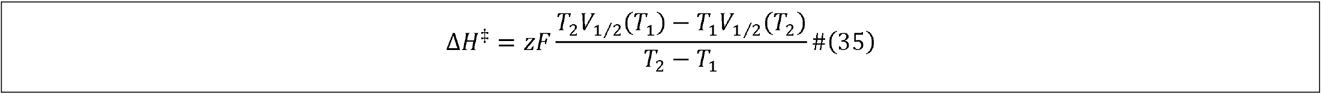

### Derivation of Δ*H*_*α*_ and Δ*H*_*β*_

We found that these magnitudes can be derived from the maximum of the time constant of current relaxation, which is given by:

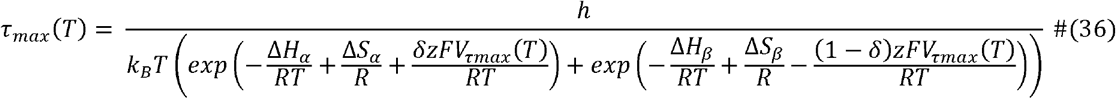

From Eq. 27 one can find *V*_*τmax*_ as:

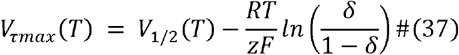

Substituting Eq. 37 in 36 yields:

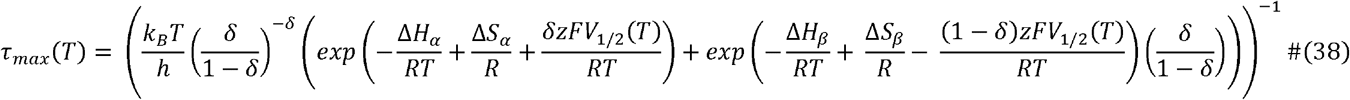

For *V* = *V*_1/2_, *α (V*_*1/2*_) is equal to *β* (*V*_1/2_), thus:

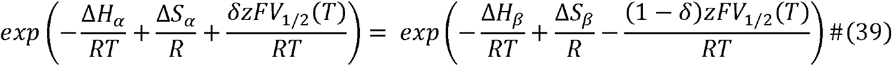

Then, taking the exponentials in Eq. 38 as common factor, *τ*_*max*_*(T)* becomes:

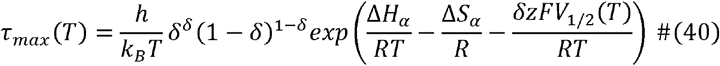

Dividing *τ*_*max*_*(T)* evaluated at two temperatures, *T*_1_ and *T*_2_:

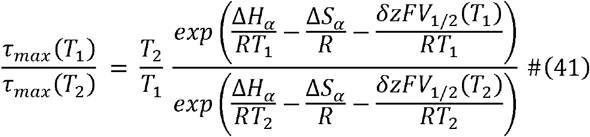

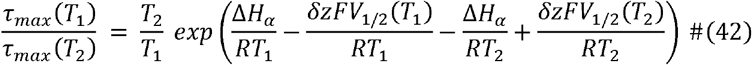

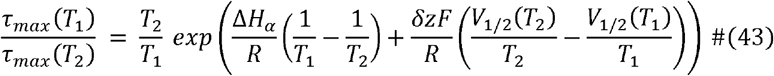

From here Δ*H*_*α*_ can be isolated as:

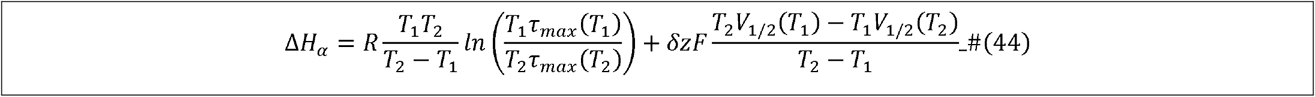

Then, Δ*H*_*β*_ can be then derived from Δ*H*_*α*_ and Δ*H*^‡^:

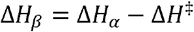

which yields:

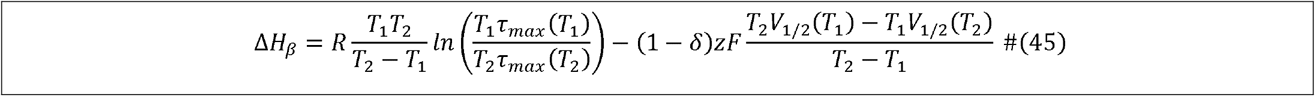

### Derivation of Δ*S*_*α*_ and Δ*S*_*β*_

Δ*S*_*α*_ can be determined by substituting Δ*H*_*α*_ from Eq. 44 in Eq. 40, evaluating in *T*_*2*_. After some algebraic calculations to isolate Δ*S*_*a*_ one obtains:

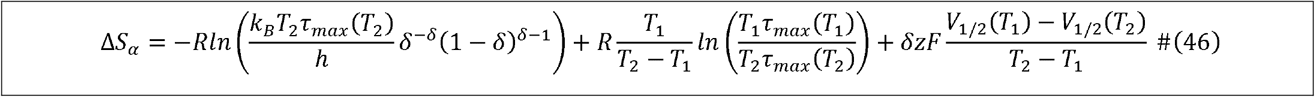

Then, Δ*S*_*β*_ can be derived from Δ*S*_*α*_ and Δ*S*^‡^:

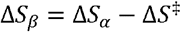

yielding:

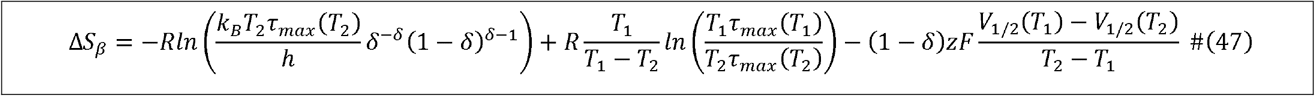

Note that for simplicity, Eq. 31, 35, 44, 45, 46 and 47 are written in terms of *z* and *δ*, which are the values calculated above with Eqs. 12 or 17 and 26 or 27, respectively.

### Practical implementation of the calculations

To illustrate the use of the derived equations in practice we employ the data published for TRPM5 (Fig. 1a, b, c). From this whole data set we selected the curves corresponding to *T*_1_= 25 °C = 298.15 K and *T*_2_ *=* 32.5 °C = 305.65 K because both of them allow the estimation of *V*_*1/2*_, the slope of *P*_*o*_ *(V, T)* evaluated in *V*_1/2_, *V*_*τ*_*max, P*_*o*_*(V, T)* evaluated in *V*_*τmax*_ and *τ*_*max*_, which are all the values required to calculate the model parameters. The visual inspection of Fig. 1d yields that *V*_1/2_(*T*_1_)≈ 160 *mV* and *V*_1/2_*(T*_2_) ≈ 105 *mV*. To determine *z* we can use Eq. 12 approximating the derivative to the slope of the data corresponding to *T*_2_ = 32.5 °C around *V*_1/2_(*T*_2_)≈ 105 *mV*. That yields:

**Fig. 1.**
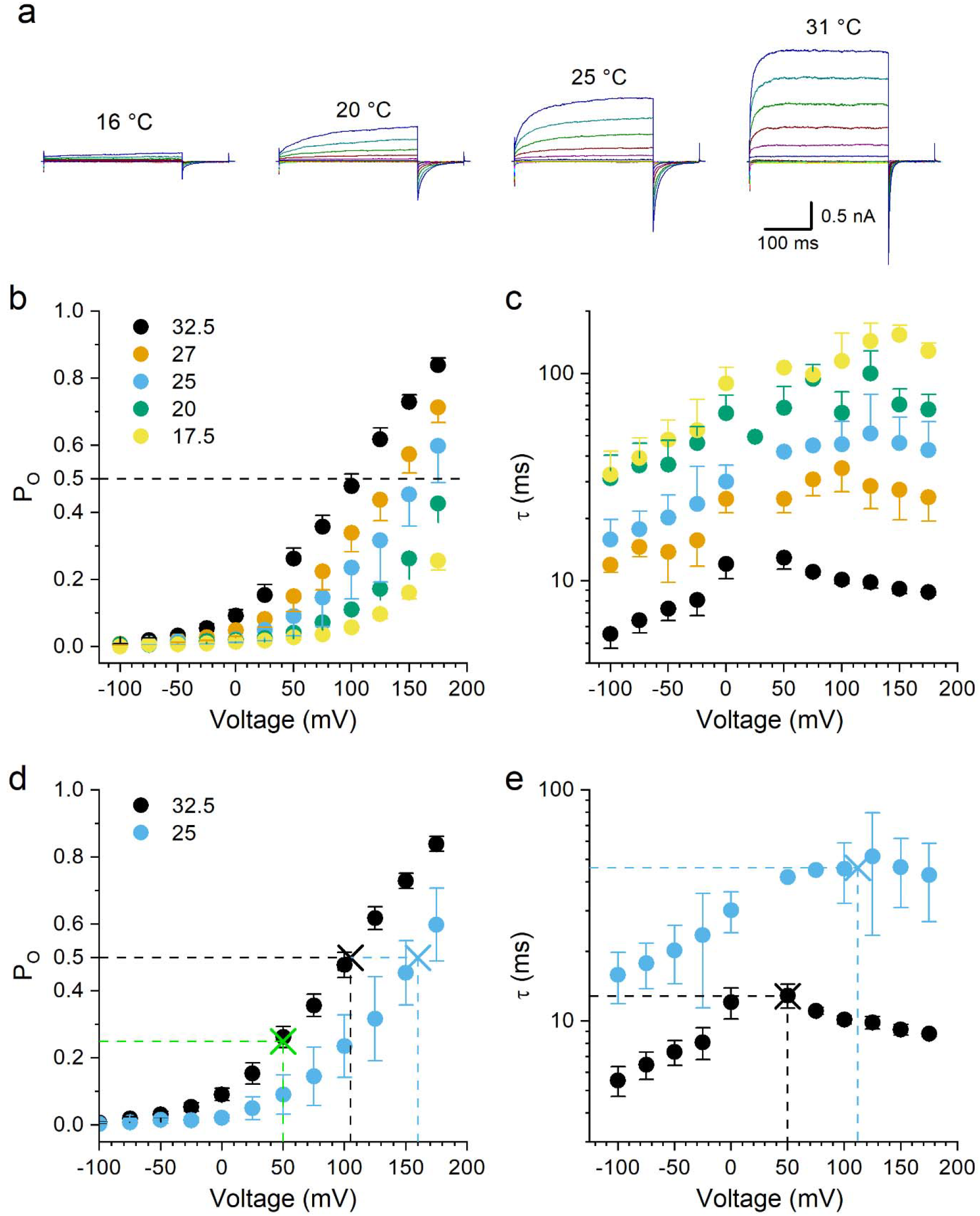
Temperature dependence of TRPM5. **a** Representative examples of TRPM5 current traces recorded in HEK293 cells at different temperatures. The currents were elicited by 200 ms-long voltage pulses given between -150 mV to +175 mV from a holding potential of +25 mV. Tail currents were elicited by a subsequent repolarizing pulse to -125 mV during 100 ms. **b, c** Average voltage dependences of the open probability and of the time constant of current relaxation at different temperatures, **d** The average voltage dependences of the open probability obtained at 32.5 and 25 °C were selected to estimate two corresponding values of voltage (*V*_1/2_) for which the activation is half-maximal (see the corresponding black and blue crosses), e The average voltage dependency of the time constant of current relaxation obtained at 32.5 and 25 °C were selected to estimate two corresponding values of maximal time constant (*τ*_*max*_, see the corresponding black and blue crosses) and the corresponding voltage values (*V*_*τmax*_). The value of *V*_*τmax*_ at 32.5 °C was also used to determine the open probability in panel **d** (green cross). The experimental data is reused from Talavera et al., [21],

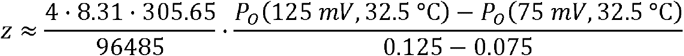

which when introducing the values of *P*_*o*_ in Fig. 1d gives:

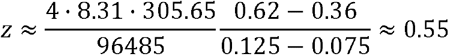

We can determine *δ* with Eq. 26, using an estimate of *V*_*τ max*_ of 50 mA/ obtained from the data at 32.5 °C in Fig. 1e and reading the value of *P*_*o*_(50 mV,32.5 °C) ≈0.25 in Fig. 1d. This yields that *δ* ≈1−0.25 = 0.75.

We can also use Eq. 27 substituting the values of *V*_*τmax*_ and *V*_1/2_ estimated at 25 °C:

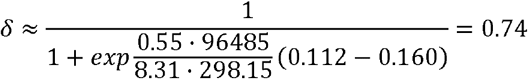

The same value can be determined using the same procedure on the data obtained at 32.5 °C:

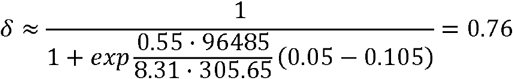

Thus, we select the value *δ* ≈ 0.75.

Δ*S*^‡^ and Δ*H*^‡^ are calculated with Eqs. 31 and 35:

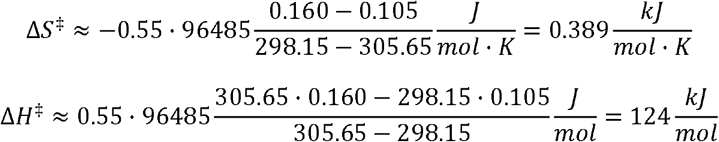

Δ*H*_*α*_ is calculated with Eq. 44 using estimates of *τ*_*max*_ found in Fig. 1e (46 ms at 25 °C and 12.8 ms at 32.5 °C):

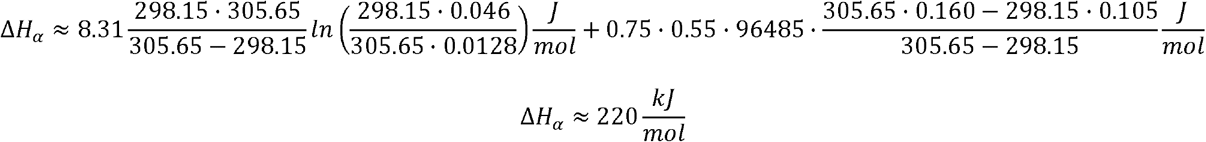

and Δ*H*_*β*_ with Eq. 45:

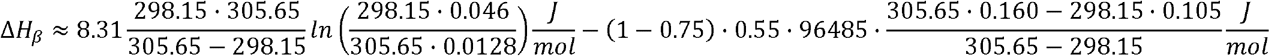

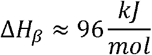

or much easier by Δ*H*_*β*_ = Δ*H* ^‡^ — Δ*H*_*α*_, which gives the same result.

Δ*S*_*a*_ is calculated with Eq. 46:

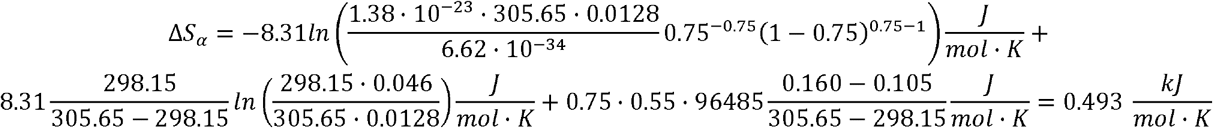

and Δ*S*_*β*_ with Eq. 47:

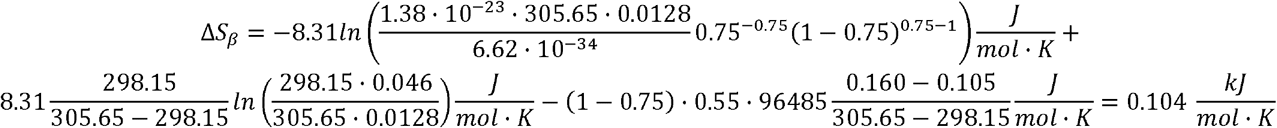

or by Δ*3*_*β*_ *=* Δ*S* ^*‡*^ *—* Δ*S*_*α*_, giving the same result.

### Global fitting

Next, we used these calculated values as initial parameters of the global fitting of the whole data set available in Fig. 1b, c. For this, we first fitted all *P*_*o*_*(V)* curves obtained at different temperatures (Fig. 2a) with the equation:

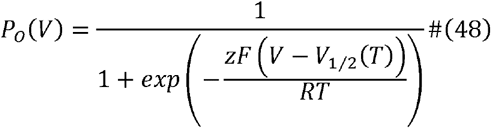

to obtain the values of *V*_1/2_ at the different temperatures and the value of z, which was considered temperature-independent. The values of *V*_1/2_ where then plotted as a function of temperature (Fig. 2b) and fitted with the linear equation (Eq. 15):

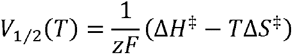

from which Δ*H* ^‡^and Δ*S* ^‡^ were obtained from the Y-axis intercept and the slope.

**Fig. 2.**
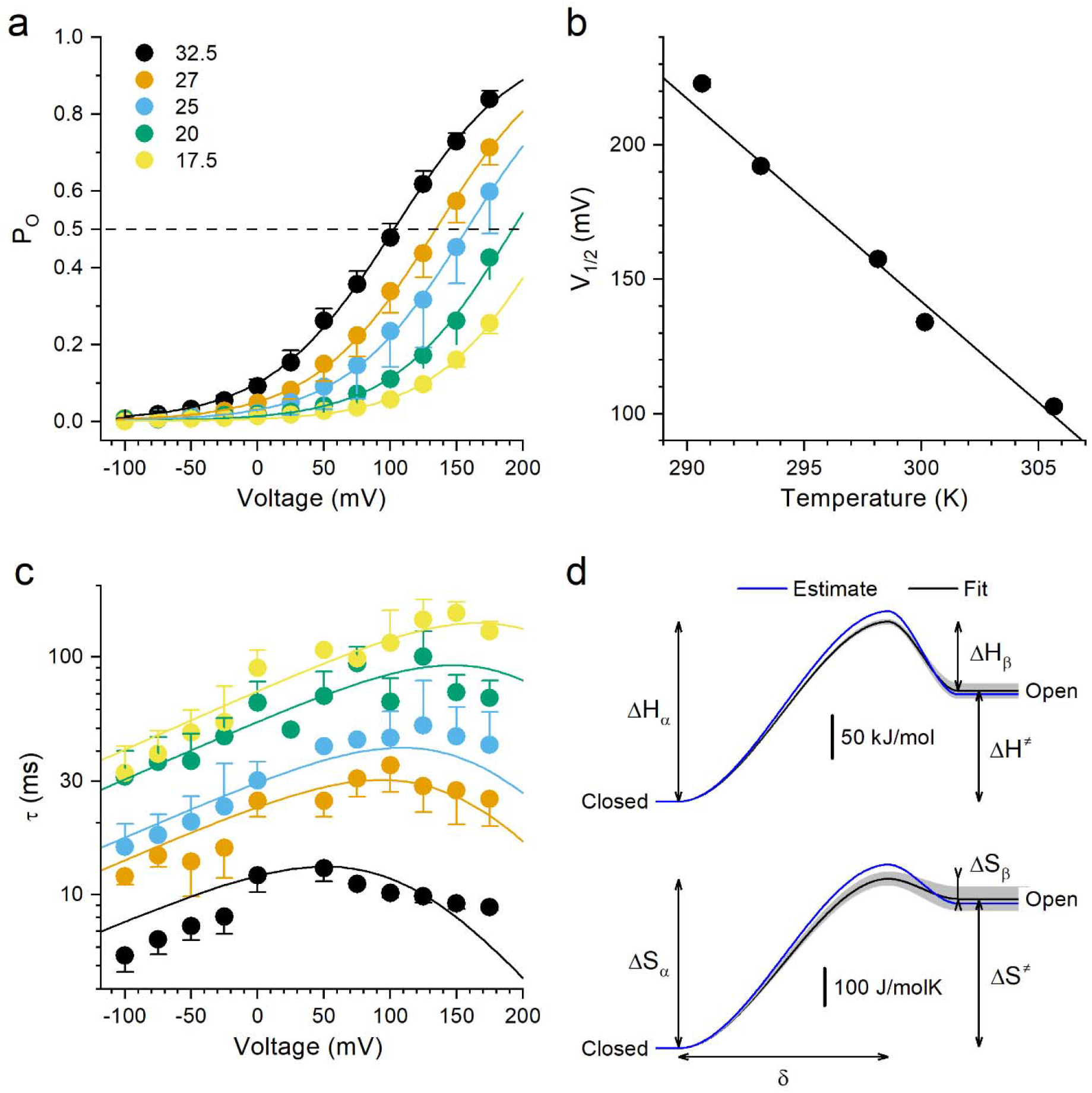
Global fit of the two-state close-open thermodynamic model to TRPM5 thermal response, **a** Global fit of the voltage dependence of the open probability using Eq. 48 to determine *V*_1/2_ for each temperature and the apparent gating charge *z*. **b** Linear fit of the temperature dependence of *V*_1/2_ with Eq. 15, from which Δ*H*^*‡*^ and Δ*S*^*‡*^ were determined from the slope and Y-axis intercept, respectively, **c** Global fit of the voltage dependence of the time constant of current relaxation at several temperatures with Eq. 49, from which Δ*H*_*α*_, Δ*S*_*α*_ and *δ* were obtained, **d** Enthalpy and entropy profiles of the model of TRPM5 drawn with the parameters estimated with the analytical equations 26, 31, 35, 44, 45, 46 and 47 or with the parameters resulting from the global fitting of the data shown in panels **a-c**. The gray bands indicate the fitting errors of the global fit parameters. The double headed arrows indicate the corresponding enthalpy and entropy values of the TRPM5 model. The experimental data is reused from Talavera et al., [21].

Then the *τ*(*V*) curves obtained at different temperatures (Fig. 2c) were fitted by a modified form of Eq. 6 in which Δ*H*_*β*_ is written as Δ*H*^‡^ — Δ*H*_*α*_ and Δ*K*_*β*_ as Δ*3*^‡^ *—* Δ.*S*_*α*_:

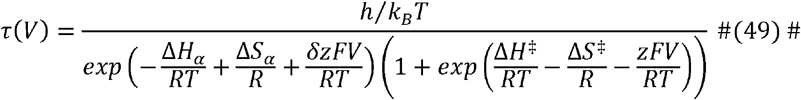

from which we obtained Δ*H*_*α*_, ΔS_*α*_ and *Δ*.

Finally, Δ*H*_*β*_ and Δ*S*_*β*_ were calculated as indicated above.

The values of most estimated parameters fell within 10% of those obtained through the global data fitting, and only the estimates for Δ*H*_*β*_ and Δ*S*_*β*_ exhibited more pronounced deviations from their final fitted counterparts (Fig. 2d and Table 1).

**Table 1.**
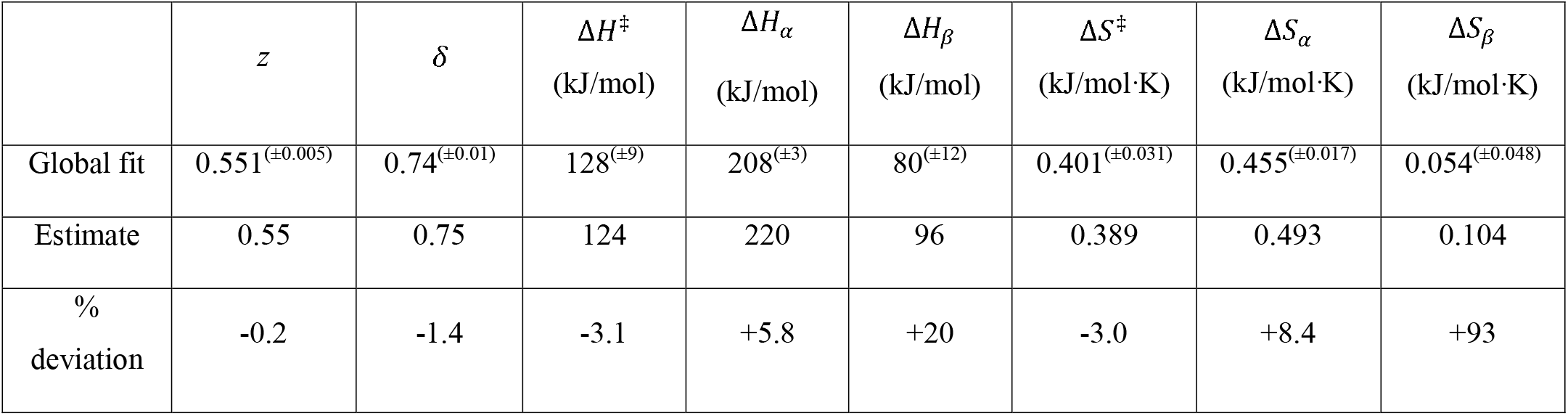
Comparison of the parameter values obtained through the global fitting procedure to those estimated using Eqs. 12 for z, 26 and 27 for an average of *δ*, and 31, 35, 44, 45, 46 and 47 for Δ*H*^*‡*^, Δ*S*^*‡*^, Δ*H*_*α*_,, Δ*S*_*α*_ ,Δ*H*_*β*_, Δ*S*_*α*_ and Δ*S*_*β*_, respectively. The superscripts in parenthesis indicate the fitting errors and the % values the relative deviation of the estimated parameters from the parameters resulting from the global fit 100%-(Estimate -Global fit)/Global fit.

## Discussion

The function of thermoTRP channels underlie one of the main molecular mechanisms by which cells detect and respond to temperature changes. Understanding their thermodynamic properties is therefore essential for elucidating the physical basis of temperature sensing and signaling in biological membranes. Parameters such as enthalpy, entropy, and heat capacity changes provide quantitative insight into the energetic landscape underlying channel activation, revealing how relatively small variations in temperature can produce large conformational transitions and steep activation profiles. Thermodynamic analyses also help distinguish between competing mechanistic models of thermo-gating, including allosteric coupling, heat-capacity driven mechanisms, and lipid-protein interactions. ThermoTRP channels are also implicated in pathological processes such as chronic pain, inflammation, and metabolic disorders. Thus, the development of robust and accessible methods to estimate thermodynamic parameters can facilitate comparative analyses across channel isoforms, experimental conditions, and species, thereby contributing to the understanding of thermoTRP channel function, regulation and the development of effective pharmacological targeting.

Here we addressed one of the main challenges in fitting thermodynamic models to thermoTRP channel data: determining an appropriate initial set of model parameters for the iterative optimization procedure. Because these models are typically nonlinear and contain multiple parameters, the fitting process can be highly sensitive to the initial selection, frequently leading to slow convergence, parameter non-identifiability, or convergence toward non-physiological parameters. We reasoned that an analytical approach for estimating thermodynamic parameters directly from salient experimental observables could provide a valuable practical solution. We selected the two-state model to implement the required algebraic calculations for being the simplest and the most widely applied to thermoTRPs [24,21,14,27], We were able to derive analytical equations for all model parameters from salient features of data acquired at only two temperatures. This is particularly advantageous in electrophysiological experiments where extensive temperature sampling may be limited by recording stability, or experimental throughput, see for instance the case for TRPA1 [14].

The equations for *δ* revealed two relations, one with the probability of the channel to be in the closed state (Eq. 26) and another with the difference between the voltage of half-maximal activation and the voltage for maximal time constant of current relaxation (Eq. 27). While in previous studies *δ* has been determined through global fitting procedures, these relations provide straightforward methods for determining this parameter directly from the graphs of *P*_*O*_(*V,T*) and *τ(V,T)*. Eqs. 31 and 35 have been previously shown to serve to determine for Δ*S*^‡^ and Δ*H*^‡^ from the slope and Y-axis intercept of the linear relationship between the *V*_1/2_ and temperature, respectively [21]. In contrast, Eqs. 44-47 for Δ*H*_*α*_, Δ*H*_*β*_, Δ*S*_*α*_ and Δ*S*_*β*_ have much more complex structures, whose geometrical interpretations deserve to be addressed in a separate study.

The implementation of the parameter estimations on data previously gathered for TRPM5 [21] yielded satisfactory results, with most estimates laying within 10% offset of the parameter values determined with the data global fitting, thereby ensuring proper convergence during the iterations. We did find stronger deviation of the estimates for two parameters (Δ*H*_*β*_ and Δ*S*_*β*_) from the final fitted values. This probably resulted from the higher dispersion of the experimental data at negative potentials, which are largely determined precisely by the closing reaction rate constant, *β*. In addition, using only data recorded at two different temperatures can limit the proper determination of the parameters. This can be remedied by, for instance, using different pairs of temperatures to estimate the parameters with the method we describe here and to average the results. There are probably other ways to obtain estimates for the model parameters using similar or other approaches, as illustrated by the fact that we found two methods (Eqs. 26 and 27) to determine estimates of *δ*. Moreover, we envisage that one can devise new methods exploiting the availability of the same salient data features (e.g., *V*_1/2_, *τ*_*max*_ and *V*_*τmax*_) at several temperatures to obtain even better parameter estimates.

An important consideration of the present analytical approach is that it is derived under the assumption that thermoTRP channel gating can be adequately described by a simple two-state closed-open model. Although this approximation has proven useful for describing the macroscopic behavior of several thermoTRP channels [24,21,14,27], it inevitably represents a simplification of the more complex conformational landscape that likely underlies thermo-gating. In reality, thermoTRP channels may transition through multiple intermediate, inactivated, sensitized, and/or ligand-dependent states, with complex coupling between temperature, voltage, and chemical stimuli. Consequently, the applicability of the present method depends on the extent to which the experimental data can be reasonably approximated by an effective two-state process. Nevertheless, the analytical strategy developed here is a conceptual foundation for extension toward more elaborate gating models. In principle, analogous derivations could be formulated for multi-state models by identifying reduced sets of experimentally measurable descriptors that constrain subsets of the thermodynamic parameters. While the resulting expressions are expected to be mathematically more complex and may no longer yield fully closed-form solutions, simplified analytical approximations could still provide valuable initial parameter estimates for subsequent global optimization procedures. Therefore, although the current method is intentionally formulated within a minimal thermodynamic framework, it may represent a first step toward the development of broader analytical tools for quantitatively characterizing complex thermoTRP gating mechanisms.

An additional future application of the present analytical formulation is the implementation of quantitative analyses of parameter sensitivity and uncertainty. In contrast to purely numerical fitting approaches, in which the relationship between experimental and parameter uncertainties may remain difficult to interpret, the analytical expressions derived here make it possible to directly examine how errors in *V*_*1/2*_, *τ*_*max*_ and *V*_*τmax*_ propagate to the estimated thermodynamic parameters. Such analyses could identify which experimental observables exert the greatest influence on the accuracy of the parameter estimates, thereby revealing potential sources of parameter uncertainty. On one hand, this could provide practical guidance about the experimental precision/accuracy required for reliable thermodynamic characterization of thermoTRP channels and help optimize experimental design by identifying the observables that most strongly constrain the model. Moreover, such analyses may serve to determine confidence intervals and robustness criteria for thermodynamic parameter estimation.

Finally, although the present study focuses on thermoTRP channel activation, the analytical strategy developed here may also be applicable to other ion channel processes that can be described using Eyring-based thermodynamic formalisms, including ion permeation voltage-dependent open pore block and mechano- and ligand-dependent gating. In these systems, experimentally measurable observables such as dose-response relationships, half-maximal agonist/inhibitory concentrations, voltage dependence of block, and reversal potentials are linked to underlying energetic parameters through nonlinear relationships similar to those analyzed here. Although extension to these applications would likely require adaptation to more elaborate multi-state schemes, the general principle of extracting energetic information directly from experimentally accessible observables may provide a useful framework for the quantitative analysis of diverse ion channel permeation and gating mechanisms.

In conclusion, we here provide analytical estimates of thermoTRP thermodynamic parameters serving as robust starting values for subsequent nonlinear fitting procedures, which may improve convergence speed, reproducibility, and fitting stability. Collectively, these advantages make direct analytical estimation a useful complement to conventional global optimization strategies for the quantitative analysis and biophysical interpretation of thermoTRP channel thermodynamics, with possible applicability to more complex gating models and to the study of permeation, block and other ion channel gating mechanisms.

## Statements and Declarations

The authors have no relevant financial or non-financial interests to disclose.

## Acknowledgements

We thank Dr. Enrique Velasco for the helpful discussions. This work was supported by grants from the Research Foundation – Flanders (FWO, G089423N) and from the Research Council of the KU Leuven (C14/23/134) to KT.

## Data availability statement

Data sets generated during the current study are available from the corresponding author on reasonable request.

## Author contributions statement

Conceptualization, M.S., M.M., I.Z-D and K.T.; investigation, M.S., M.M., I.Z-D and K.T.; writing—original draft preparation, M.S., M.M., I.Z-D and K.T.; writing—review and editing, M.S., M.M., I.Z-D and K.T.; visualization, M.S. and K.T.; supervision, K.T.; project administration, K.T.; funding acquisition, K.T. All authors have read and agreed to the published version of the manuscript.

## Notes

### Competing Interest Statement

The authors have declared no competing interest.

## References

1. Brauchi S, Orio P, Latorre R (2004) Clues to understanding cold sensation: thermodynamics and electrophysiological analysis of the cold receptor TRPM8. Proc Natl Acad Sci U S A 101:15494–15499. doi: 10.1073/pnas.0406773101

2. Brauchi S, Orta G, Mascayano C, Salazar M, Raddatz N, Urbina H, Rosenmann E, Gonzalez-Nilo F, Latorre R (2007) Dissection of the components for PIP2 activation and thermosensation in TRP channels. Proc Natl Acad Sci U S A 104:10246–10251. doi: 10.1073/pnas.0703420104

3. Brauchi S, Orta G, Salazar M, Rosenmann E, Latorre R (2006) A hot-sensing cold receptor: C-terminal domain determines thermosensation in transient receptor potential channels. J Neurosci 26:4835–4840. doi: 10.1523/JNEUROSCI.5080-05.2006

4. Clapham DE, Miller C (2011) A thermodynamic framework for understanding temperature sensing by transient receptor potential (TRP) channels. Proc Natl Acad Sci U S A 108:19492–19497. doi: 10.1073/pnas.1117485108

5. Cordero-Morales JF, Gracheva EO, Julius D (2011) Cytoplasmic ankyrin repeats of transient receptor potential A1 (TRPA1) dictate sensitivity to thermal and chemical stimuli.Proc Natl Acad Sci U S A 108:E1184–1191.doi: 10.1073/pnas.1114124108

6. Diaz-Franulic I, Verdugo C, Gonzalez F, Gonzalez-Nilo F, Latorre R (2021) Thermodynamic and structural basis of temperature-dependent gating in TRP channels.Biochem Soc Trans 49:2211–2219.doi:10.1042/BST20210301

7. Femandez-Ballester G, Fernandez-Carvajal A, Ferrer-Montiel A (2023) Progress in the Structural Basis of thermoTRP Channel Polymodal Gating.Int J Mol Sci 24.doi:10.3390/ijms24010743

8. Garcia-Elias A, Mrkonjic S, Pardo-Pastor C, Inada H, Hellmich UA, Rubio-Moscardo F, Plata C, Gaudet R, Vicente R, Valverde MA (2013) Phosphatidylinositol-4,5-biphosphate-dependent rearrangement of TRPV4 cytosolic tails enables channel activation by physiological stimuli.Proc Natl Acad Sci U S A 110:9553–9558.doi:10.1073/pnas.1220231110

9. Grandi J, Hu H, Bandell M, Bursulaya B, Schmidt M, Petrus M, Patapoutian A (2008) Pore region of TRPV3 ion channel is specifically required for heat activation.NatNeurosci 11:1007–1013.doi:10.1038/nn.2169

10. Grandi J, Kim SE, Uzzell V, Bursulaya B, Petrus M, Bandell M, Patapoutian A (2010) Temperature-induced opening of TRPV1 ion channel is stabilized by the pore domain.NatNeurosci 13:708–714.doi:10.1038/nn.2552

11. Hille B (2001) Ionic Channels of Excitable Membranes.3rd edn.Sinauer.

12. Hodgkin AL, Huxley AF (1952) A quantitative description of membrane current and its application to conduction and excitation in nerve.J Physiol 117:500–544.doi:10.1113/jphysiol.1952.sp004764

13. Jara-Oseguera A, Islas LD (2013) The role of allosteric coupling on thermal activation of thermo-TRP channels.Biophys J 104:2160–2169.doi:10.1016/j.bpj.2013.03.055

14. Karashima Y, Talavera K, Everaerts W, Janssens A, Kwan KY, Vennekens R, Nilius B, Voets T (2009) TRPA1 acts as a cold sensor in vitro and in vivo.Proc Natl Acad Sci U SA 106:1273–1278.doi:10.1073/pnas.0808487106

15. Kashio M, Tominaga M (2022) TRP channels in thermosensation.Curr Opin Neurobiol 75:102591.doi: 10.1016/j.conb.2022.102591

16. Nilius B, Flockerzi V (2014) Mammalian transient receptor potential (TRP) cation channels.Preface.Handb Exp Pharmacol 223:v–vi

17. Patapoutian A, Peier AM, Story GM, Viswanath λ (2003) ThermoTRP channels and beyond: mechanisms of temperature sensation.Nat Rev Neurosci 4:529–539.doi:10.1038/nrnll41

18. Pertusa M, Gonzalez A, Hardy P, Madrid R, Viana F (2014) Bidirectional modulation of thermal and chemical sensitivity of TRPM8 channels by the initial region of the N-terminal domain.J Biol Chem 289:21828–21843.doi: 10.1074/jbc.Ml14.565994

19. Reeh PW, Fischer MJM (2022) Nobel somatosensations and pain.Pflugers Arch 474:405–420.doi:10.1007/s00424-022-02667-x

20. Talavera K, Nilius B (2011) Electrophysiological Methods for the Study of TRP Channels.In: Zhu MX (ed) TRP Channels. Boca Raton (FL),

21. Talavera K, Yasumatsu K, Voets T, Droogmans G, Shigemura N, Ninomiya Y, Margolskee RF, Nilius B (2005) Heat activation of TRPM5 underlies thermal sensitivity of sweet taste.Nature 438:1022–1025.doi:10.1038/nature04248

22. Vlachova V, Teisinger J, Susankova K, Lyfenko A, Ettrich R, Vyklicky L (2003) Functional role of C-terminal cytoplasmic tail ofratvamlloidreceptor 1.JNeurosci23:1340–1350.doi: 10.1523/JNEUROSCI.23-04-01340.2003

23. Voets T (2012) Quantifying and modeling the temperature-dependent gating of TRP channels.Rev Physiol Biochem Pharmacol 162:91–119.doi: 10.1007/112_2011_5

24. Voets T, Droogmans G, Wissenbach U, Janssens A, Flockerzi V, Nilius B (2004) The principle of temperature-dependent gating in cold- and heat-sensitive TRP channels.Nature 430:748–754.doi:10.1038/nature02732

25. Voets T, Owsianik G, Janssens A, Talavera K, Nilius B (2007) TRPM8 voltage sensor mutants reveal a mechanism for integrating thermal and chemical stimuli.Nat Chem Biol 3:174–182.doi: 10.1038/nchembio862

26. Vriens J, Nilius B, Voets T (2014) Peripheral thermosensation in mammals.Nat Rev Neurosci 15:573–589.doi:10.1038/nm3784

27. Vriens J, Owsianik G, Hofmann T, Philipp SE, Stab J, Chen X, Benoit M, Xue F, Janssens A, Kerselaers S, Oberwinkler J, Vennekens R, Gudermann T, Nilius B, Voets T (2011) TRPM3 is a nociceptor channel involved in the detection of noxious heat.Neuron 70:482–494.doi: 10.1016/j.neuron.2011.02.051

28. Yang F, Cui Y, Wang K, Zheng J (2010) Thermosensitive TRP channel pore turret is part of the temperature activation pathway.Proc Natl Acad Sci U S A 107:7083–7088.doi: 10.1073/pnas.1000357107

29. Yao J, Liu B, Qin F (2011) Modular thermal sensors in temperature-gated transient receptor potential (TRP) channels.Proc Natl Acad Sci U S A 108:11109–11114.doi : 10.1073/pnas.1105196108

30. Yeh F, Jara-Oseguera A, Aldrich RW (2023) Implications of a temperature-dependent heat capacity for temperature-gated ion channels.Proc Natl Acad Sci U S A 120:e2301528120.doi:10.1073/pnas.2301528120

